# *Dnmt3a* mutations limit normal and autoreactive Tfh differentiation

**DOI:** 10.1101/2024.02.16.580463

**Authors:** Yunbing Shen, Zhaojun Li, Sanjaykumar Boddul, Zsolt Kasza, Alexander Espinosa, Lars Klareskog, Fredrik Wermeling

## Abstract

Rheumatoid arthritis (RA) is an autoimmune disease characterized by joint inflammation, strongly associated with the activity of autoreactive CD4+ T cells. *DNMT3A* mutations are the most common somatic mutations found in the hematopoietic system of patients with rheumatoid arthritis. However, the role of DNMT3A in CD4+ T cells and CD4+ T follicular helper (Tfh) cells is poorly understood. Since somatic mutations are not identified in standard genome-wide association studies, somatic mutations’ impact on the etiology of diseases could be underestimated. Here, we thoroughly characterized and used the KRN+ splenocyte transfer model of autoimmune joint inflammation and inactivated *Dnmt3a* using CRISPR-Cas9 and standard Cre/loxP approaches. Experiments with competitive bone marrow (BM) chimeras identified a positive role for *Dnmt3a* in Tfh differentiation, which was validated by comparing mice with *Dnmt3a* mutations in CD4+ cells to animals with WT *Dnmt3a*. In conclusion, We identify that *Dnmt3a* mutations limit normal and autoreactive Tfh differentiation.

**Key findings:**

– *Dnmt3a* mutations limit Tfh differentiation, which could contribute to reduced immune responses in individuals with somatic *DNMT3A* mutations.
– Deep characterization of the KRN+ splenocyte transfer model defines a dynamic process leading to reproducible autoimmune joint inflammation.
– The immuno-CRISPR (iCR) methodology can be used to test the role of candidate genes in disease models.

## 1. Introduction

Rheumatoid arthritis affects approximately 0.5% of the global population with a female bias (*1*). Genome-wide association studies (GWAS) have associated specific HLA-DR MHC class II alleles with an increased risk of developing RA (particularly with the seropositive subgroup, characterized by patients with a spectrum of autoantibodies and a worse prognosis) (*2*). This observation connects MHC class II antigen presentation and activation of CD4+ T cells to the disease etiology and progression (*2-4*). Multiple minor GWAS hits have also associated CD4+ T cells with the disease by mapping to important functions in CD4+ T cells (*5, 6*).

KRN+ TCR transgenic animal models for autoimmune arthritis are based on KRN+ CD4+ T cells recognizing an autoantigen presented in I-A^g7^ MHC class II (*7*). This results in the production of arthritogenic autoantibodies (anti-GPI) and erosive arthritis (*8*). Models based on KRN+ TCR transgenic mice, thus, share features of seropositive RA (a link to MHC class II presentation and autoantibody production), and could be used as a preclinical *in vivo* model to study autoreactive CD4+ T cells in the context of autoimmune joint inflammation. “KRN” is the name of the specific TCR used to make the KRN TCR transgenic animals.

Somatic mutations are mutations that accumulate in cells over an individual’s lifetime due to DNA’s inherent instability (*9*). Such mutations are central to cancer development, but an increasing body of work has identified that somatic mutations also could play a role in other diseases (*10*). Clonal hematopoiesis is the condition where specific somatic mutations in hematopoietic stem cells (HSCs) result in a selective advantage of the modified HSCs clone and, over time, an increased contribution to hematopoiesis by the expanded HSCs clone (*11*). Consequently, the specific HSC mutations become increasingly found in mature immune cells over time and could theoretically affect their function and selection. It has been reported that more than 60% of individuals >80 years show evidence of clonal hematopoiesis, and 20-30% of 50-60-year-olds (*12*). Other studies have reported more modest frequencies (*13*) but also higher frequencies (14), likely depending on the sensitivity of the method used. Most studies start identifying evidence for clonal hematopoiesis in individuals from 30 years of age. However, one study reported that clonal hematopoiesis was found in 3% of 20-29-year-old individuals using a high-sensitivity method (*14*). Importantly, the frequency of immune cells with the clonal hematopoiesis-linked mutation is expected to be low, and therefore, standard bulk sequencing efforts will not identify such mutations. GWAS studies are, thus, likely not to identify somatic mutations, even if the source material for the sequencing is blood. Therefore, the contribution of mutations in clonal hematopoiesis genes to immune-related aberrations and diseases could be underestimated.

The most commonly mutated genes in clonal hematopoiesis are *DNMT3A* and *TET2*, two genes involved in DNA methylation (*11*). Due to the role of DNA methylation, mutations in these two genes are expected to broadly affect many different pathways (*15*). Of note, one study of RA patients identified *DNMT3A* mutations as the strongest contributor to clonal hematopoiesis in the studied group (*16*).

## 2. Material and Methods

Primers, reagents, sourcing, and catalog numbers can be found in Tables S1-4.

### 2.1. Animals

8 to 12 weeks old, age and sex-matched animals were used for experiments. All animal experiments were approved by the ethical board at Karolinska Institutet. All mice were housed in SPF conditions in a 12-hour light and 12-hour dark cycle and fed with a standard chow diet *ad libitum*. The B6.KRN mouse strain was a kind gift from Diane Mathis and Christophe Benoist (Harvard Medical School) (*7*). Cas9 GFP mouse strain (stock no. 026179), CD45.1 mouse strain (stock no. 002014), TCRb KO mouse strain (stock no. 002118), NOD mouse strain (stock no. 001976), CD4^cre^ mouse strain (stock no. 022071), Dnmt3^fl-R878H^ (stock no. 032289) were purchased from Jackson Laboratory. KRN Cas9.1 mouse strain was generated by crossing B6.KRN, Cas9 GFP, and CD45.1 mouse strains. *TCRb*^*-/-*^ *I-A*^*b*^ *I-A*^*g7*^ mouse strain was generated by crossing TCRb KO and NOD mouse strains. CD4^cre^ Dnmt3a^fl/fl^ mouse strain was generated by crossing *Cd4*^*cre*^ and Dnmt3a^fl-R878H^ mouse strains. Genotyping primers are listed in Table S1.

### 2.2. CRISPR-modifying BM cells and BM transplantation

BM cells were prepared as described before (*17*). For BM transplantation, roughly 10^6^ BM cells were i.v. injected into recipient mice lethally irradiated (900 rad) 6-18h previously. For mixed BM chimera experiment, lineage-depleted BM cells were i.v. injected into lethally irradiated recipient mice at a 1:1 ratio (roughly 5×10^5^ of each genotype). Subsequent experiments were conducted at least 8 weeks after BM transplantation.

### 2.3. KRN T cell transfer

KRN+ splenocytes were prepared by pressing the spleen through a 40 mm cell strainer with a 3 mL syringe plunger. Roughly 2*10^7^ cells were injected i.v. via the tail vein into *TCRb*^*-/-*^*I-A*^*b*^*/I-A*^*g7*^ recipient mice. The severity of arthritis was scored every two to three days by clinical examination of each paw and ankle (0, no swelling; 3, maximal swelling), adding up to a total clinical score (0–12) per mouse, like in (*18, 19*). The weight of the animals was monitored every two to three days. At the end of each experiment, organs and blood were collected for further analysis.

### 2.4. B cell depletion in KRN T cell transfer model

75 ug/mouse/injection of the anti-CD20 antibody (Bioxcell, Cat. No. BE0356) was i.v. injected before (day -3 and -1), and after (days 1 and 3), the KRN T cell transfer. Control mice were injected with vehicle. The weight and clinical score were examined as indicated in the figure. At the end of each experiment, organs and blood were collected for further analysis.

### 2.5. Serum transfer

Blood from KRN T cell transfer recipient mice was collected into sterile 1.5 ml Eppendorf tubes (days 9-15, depending on the experiment). Serum was prepared by allowing the blood to clot for at least 1 hour, centrifuge at 10,000 rpm for 10 minutes, followed by collection of the serum layer, subsequently stored at -20°C. Serum transfer experiments were performed with pooled serum preparations from several donors by i.v. injection of 200 ul serum into WT C57BL/6 mice. Joint inflammation and body weight were followed as indicated in the figure.

### 2.6. ELISA

IL-6 ELISA was performed according to the manufacturer’s instruction (Biolegend, Cat. No. 431301) with mouse serum diluted 1:50. Anti-GPI antibody ELISA was conducted by coating ELISA plates (Invitrogen, Cat. No. 44-2404-21) with 50 μl of 10 μg/ml Glucose-6-Phosphate Isomerase (SIGMA, Cat. No. P5381) at 4°C overnight. Wells were washed in PBS + 0.05% Tween, and after a 2-12h blocking step (PBS + 2% BSA) at RT (2h incubation) or 4°C (overnight/12h) diluted mouse sera (1:100-1:500 in block buffer) were added into the wells. After 2-12h incubation, the wells were washed, and HRP conjugated anti-mouse IgM (Santa Cruz Biotechnology, Cat. No. sc-2973) or IgG (Santa Cruz Biotechnology, Cat. No. sc-2005) diluted 1:10000 In block buffer was added. After a two-hour incubation at room temperature, the wells were washed, TMB substrate (Medicago, Cat. No. 10-9405-250) was added, and the plate was read by an ELISA reader at 450 nm.

### 2.7. SRBC immunization

Sheep red blood cells (SRBC) were purchased from Håtunalab AB (Stockholm, Sweden). 1 ml SRBCs were washed in PBS three times (1000g for 10 minutes) and resuspended in 1 ml PBS. Mice were subsequently immunized i.p. with 200 ul of the SRBC solution. For Tfh characterization, immunized mice were sacrificed on day 7, and organs were collected for further analysis. For antibody production analysis of, blood was sampled 14 days after immunization.

### 2.8. Detection of serum SRBC-specific antibodies

SRBC-specific antibodies were measured as described in (*20*). Briefly, SRBCs were diluted 1:10 in PBS, and 20 ul of the SRBC solution was mixed with 1 ul of serum. After a 20-minute incubation at 4°C, the SRBCs were washed in a standard FACS buffer (1000g for 10 minutes) and stained with anti-mouse IgG-BV605 (BD, 405327) or anti-IgM (BD, 406517) diluted 1:200 for 20 minutes at 4°C. Samples were washed with FACS buffer, resuspended in 200 ul of FACS buffer, and acquired by flow cytometry (Cytek, Aurora).

### 2.9. Flow cytometry analysis and sorting

Cell suspension from organs was prepared by pressing organ/s through a 40-micrometer cell strainer. The cells were stained with antibodies according to manufacturers’ recommendations (Table S4). Briefly, wells were stained in cold MACS buffer (Miltenyi Biotec, Cat. No.130-091-221) at 4°C for 30 minutes, washed with MACS buffer, and resuspended in MACS buffer for further analysis. Samples were acquired by BD LSR Fortessa and Cytek, Aurora, or sorted by SONY SH800. Generated FCS files were analyzed by Flowjo (V10, Tree Star Inc).

### 2.10. sgRNA, primer design, and Indel analysis

sgRNAs were designed as described previously (*17*). Briefly, the Green Listed software http://greenlisted.cmm.ki.se/ (*21*) and Synthego CRISPR design tool were utilized to design sgRNAs (Table S2). Primers for sequencing the sgRNA target sites were designed using Primer Blast, aiming for a 400-800 bp amplicon with the sgRNA binding site in the middle (Table S3). Genomic DNA from sorted cells was collected using DNeasy Blood & Tissue Kits (Qiagen, Cat. No. 69504). The sgRNA target sites were amplified using PCR and purified using the DNA clean and concentrate kit (Zymo Research, Cat. No. D4033). The purified PCR amplicon products were sequenced by Eurofins Genomics, and Indel profiles were analyzed by Synthego ICE tool (https://ice.synthego.com).

### 2.11. Statistical analysis

GraphPad Prism version 9 was used as indicated in figure legends.

## 3. RESULTS

### KRN+ splenocyte transfer results in a reproducible and timed joint inflammation

To study how autoreactive CD4+ T cells can contribute to autoimmune joint inflammation, we established the KRN+ splenocyte transfer model. In the model (**Fig. 1A**), purified T cells or splenocytes from KRN TCR transgenic C57BL/6 mice (B6.KRN) are transferred to T-cell deficient mice expressing one copy of the C57BL/6 MHC II (I-A^b^) and one copy of the NOD MHC II (I-A^g7^, responsible for presenting the arthritogenic epitopes recognized by the KRN TCR (*7*). By crossing the B6.KRN mice with B6.CD45.1 mice, the transferred cells could be tracked in the CD45.2+ recipient mice. We found that the transfer resulted in a very timed and reproducible joint inflammation with 100% penetrance and a concomitant drop in body weight, with a peak change seen from day 6 to day 9 (**Fig. 1B-C**). A kinetic study tracking the total amount of CD4+ T cells (CD45.1+) in different organs showed that the transferred T cells were readily found in all tested organs, although only minute levels were found in the blood and the Peyer’s patches, while the most dominant populations were found in spleen and mesenteric lymph nodes (LN). In many of the secondary lymphoid organs, we found a significant increase of CD4+ T cells comparing day 6 to day 3, indicating an active expansion occurring at this stage where no clinical disease was yet seen (**Fig. 1D and S1A**). An expansion of germinal center (GC) B cells mirrored the expansion of CD4+ T cells, with a peak seen at day 6, predominately found in spleen and mesenteric LN (**Fig. 1D and S1B**). As expected, the GC B cells were >95% CD45.2+, in line with the B cells of the CD45.2+ recipient mice expressing I-A^g7^ needed for the presentation of the arthritogenic peptides (**Fig. S1B**). Interestingly, looking at T- and B-cell activation based on CD69 expression, a difference in dynamic was seen, with a seemingly delayed activation seen in the T-cell compartment, with a day 12 peak of response (**Fig. S1C-D**), in comparison to the B-cell compartments with a peak seen already at day 3 (**Fig. S1E-F**). Concomitant with the dynamics of the lymphocyte populations, we found significant levels of IgM anti-GPI already at day 6 (**Fig. 1F**) and significant IgG anti-GPI levels from day 9 (**Fig. 1G**). Notably, the circulating autoantibody levels clearly dropped at day 15 despite the joint swelling and drop in body weight still remaining. In line with the transient nature of the autoantibody levels, serum levels of IL-6 peaked at day 9 and then distinctly declined (**Fig. 1H**).

**Figure 1.**
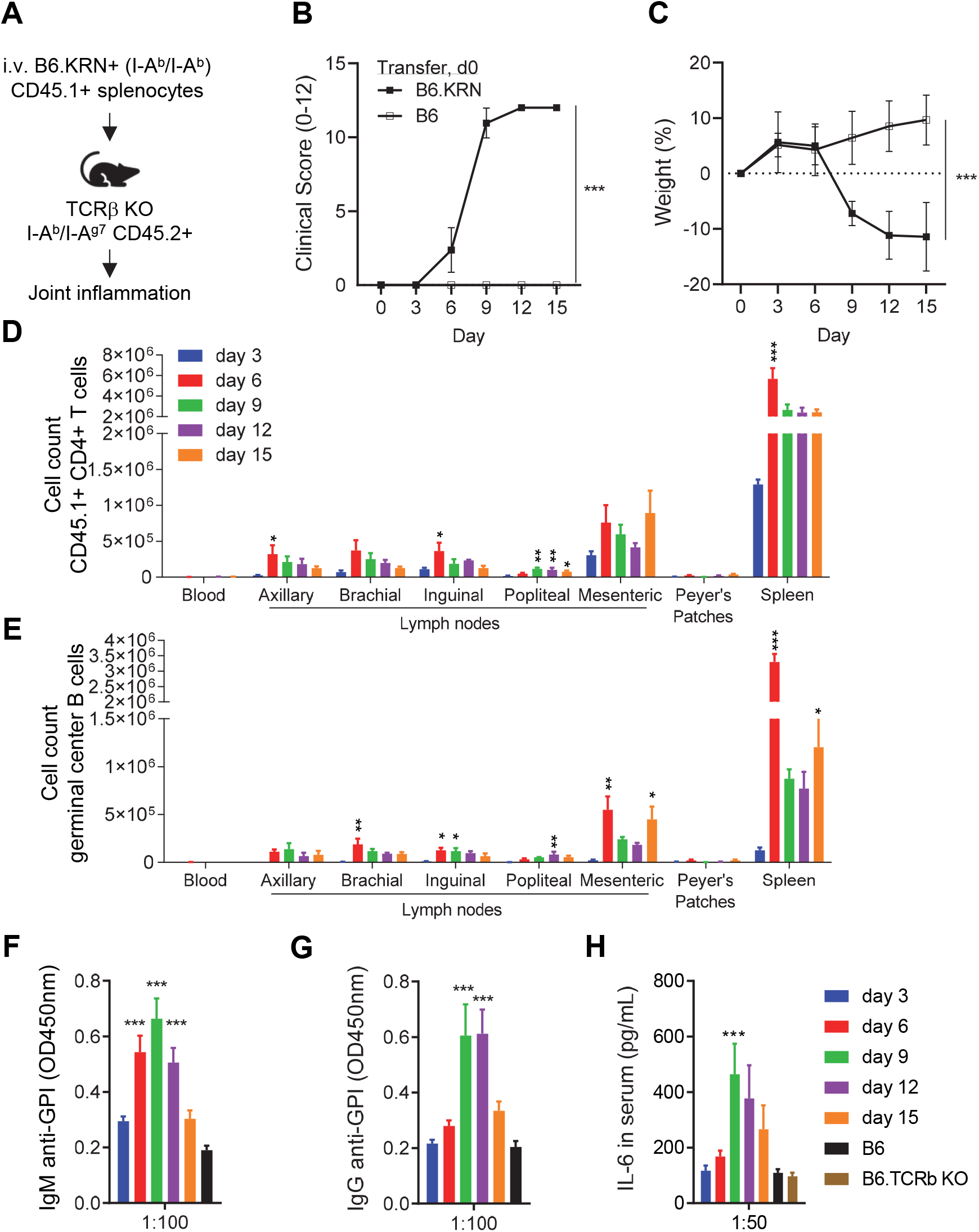
KRN+ splenocyte transfer results in a reproducible and timed joint inflammation. (**A**) Schematic describing the transfer of KRN+ splenocytes into T-cell deficient (TCRβ KO) mice expressing one copy of B6 MHCII and one copy of NOD MHCII (I-A^g7^) resulting in joint inflammation. (**B-C**) Clinical score (B), and body weight change (C) following transfer of KRN+ or control splenocytes. (**D**) Total amount of transferred CD4+ T cells (viable, singlet, CD45.1+, CD4+) detected by flow cytometry in different organs at indicated timepoints. (**E**) Germinal center (viable, singlet, CD19+, CD95+ GL7+) measured by flow cytometry in different organs at indicated time (**F-G**) Anti-GPI ELISA showing IgM (F) and IgG (G) levels at indicated time points. (**H**) IL-6 measured by at indicated time points. Data presented as mean and SEM (B-H), B-C n=20 for KRN group and 5 for ctrl; D-F-G n=3-5. *** P <0.005, ** P <0.01, and * P <0.05 by two-way ANOVA with Tukey’s post hoc test (B-C), or ay ANOVA with Dunnett’
ss post hoc test comparing different timepoints to day 3 group (D-E), or comparing different timepoints to to T-cell deficient TCRb KO group (F-H). Data is representative for two or more independent experiments.

We concluded that the KRN+ splenocyte transfer model represents a simple and robust model to study autoimmune joint inflammation. Additionally, we describe lymphocyte dynamics that complement the characterization of the model by LeBranche *et al*. (*22*).

### Joint inflammation in the KRN splenocyte transfer model is abrogated by B-cell depletion

Studies using the genetic version of the K/BxN model have identified a central pathogenic role for arthritogenic antibodies in the model (*8*). Therefore, we hypothesized that B-cell depletion in the KRN+ splenocyte transfer model would limit the resulting joint inflammation, supporting a mechanism where KRN+ T cells predominately play a pathogenic role by contributing to autoreactive germinal centers and autoantibody production. To this end, we injected mice with anti-CD20 (**Fig. 2A**) and followed how this affected the disease development. We found that the B-cell depletion completely abrogated the joint inflammation and drop in body weight found in control animals (**Fig. 2B-C**). Flow cytometry of spleens at the end of the experiment identified a strong depletion of total B cells (**Fig. 2D**) and GC B cells (**Fig. 2E**). In contrast, no difference in the levels of KRN+ T cells was found (**Fig. 2F**), nor in KRN+ T follicular helper (Tfh) cells (**Fig. 2G**). Additionally, no apparent difference in CD69 levels was found in T cells comparing control and anti-CD20 treated mice (**Fig. S2C**). The anti-CD20 injected animals showed low IgM and IgG autoantibody levels (**Fig. 2H**) as well as low IL-6 levels (**Fig. 2I**), in line with the joint inflammation and B-cell depletion phenotypes.

**Figure 2.**
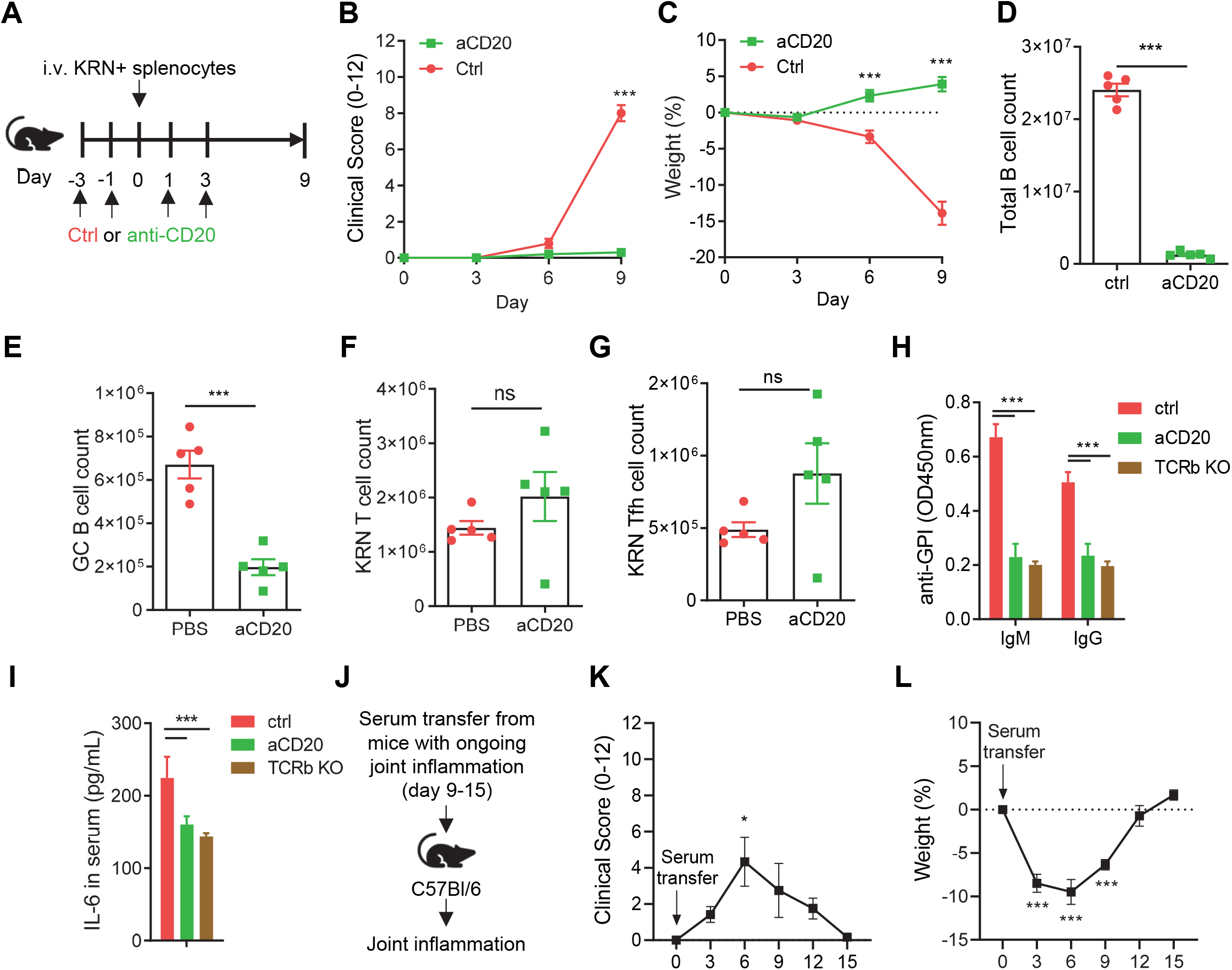
Joint inflammation in the KRN T cell transfer model is abrogated by B-cell depletion. (**A**) Schematic describing the B-cell depletion experiment. (**B-C**) Clinical score (B) and weight change (C) at indicated time points. Total number of B cells (D), germinal center B cells (E), KRN+ T cells (F), and KRN+ T follicular helper (Tfh) spleens from mice collected at day 9. (**H-I**) IgM and IgG anti-GPI levels (G), and IL-6 (H) determined by in serum collected day 9. (**J-L**) Pooled serum collected day 9-15 from mice with joint inflammation triggered by T cells transfer was i.v. injected into recipient C57BL6 mice. Clinical score (K) and weight change (L) at indicated time points. Representative flow cytometry gating is found in Fig. S2. Data presented as mean and SEM (B-I, K-L), 5; H-I n=4-5; K-L n=6. *** P <0.005, * P <0.05, and ns = non-significant by two-way ANOVA with Sídác post t (B-C), unpaired T test (D-F), or one-way ANOVA with Tukey’s post hoc test comparing samples to the time point 0 (F-H). Data is representative for two or more independent experiments.

Taken together, this implied a central role for the anti-GPI autoantibodies in the triggered joint inflammation. To test if arthritogenic autoantibodies were produced in the recipient mice, we transferred pooled serum collected from mice with ongoing joint inflammation triggered by KRN+ splenocyte transfer (**Fig. 2J**). We observed that the serum transfer triggered joint inflammation (**Fig. 2K**) and a drop in body weight (**Fig. 2L**). The joint swelling and body weight drop were both transient and noticeably milder than the mice from which the serum was collected. However, a mild phenotype could be expected, as only a fraction of the total serum volume was transferred from the affected mice to the recipients.

We concluded that the joint inflammation induced by KRN+ T cells depends on B cells, highlighting the role of autoreactive CD4+ T cells as Tfh cells promoting autoreactive germinal center reactions and subsequent autoantibody production.

### A CRISPR-based mixed bone marrow chimera identifies a role for Dnmt3a for autoreactive Tfh differentiation

As we had established parameters linked to the joint inflammation in the KRN+ splenocyte model, we next aimed to test if we could identify genes that affect the disease development (**Fig. 3A**). To do this, we resorted to an *in vivo* CRISPR-based approach that we previously published (*23*) and adopted it to the KRN+ splenocyte transfer model. By electroporating sgRNAs targeting genes of interest into Cas9+ BM cells, followed by transplanting these modified BM cells into irradiated recipients, we generate what we refer to as immuno-CRISPR (iCR) mice. As CRISPR-Cas9 generates a heterogenous gene editing result, including cells with different mutations as well as unmodified wild-type (WT) alleles, the resulting iCR mice will develop an immune system (from the transplanted CRISPR-targeted BM cells) with a controlled genetic heterogeneity at the targeted site. Next, we identify by sequencing how the genetic heterogeneity is enriched or depleted in different cell populations, different activation states, or in cells at different localizations. Essentially, this shares many features of traditional BM chimera models, where typically WT and KO BM cells are transplanted into an irradiated host, and the cell-intrinsic role of the modified gene is identified by flow cytometry (taking advantage of that the WT and KO BM typically are congenically marked with CD45.1 and CD45.2, which subsequently can be used as a proxy for the genotype of a cell). In the iCR setup, we instead sort the cell populations of interest and sequence the targeted gene. Here, we generated KRN+ iCR mice (no I-Ag7, so no disease), transferred the KRN+ iCR splenocytes to I-A^g7^+ recipients, and followed how the mutations segregated between CD4+ Tfh cells and CD4+ non-Tfh cells as the disease developed. Aiming to find genes involved in the differentiation of autoreactive Tfh cells, we first targeted *Bcl6*, necessary for Tfh differentiation (*24*) and for arthritis induced by KRN+ T cell transfer (*25*), as a positive control. Sorting splenic CD4+ Tfh cells and CD4+ non-Tfh cells from I-A^g7^+ recipients of KRN+ *Bcl6* iCR mice showed a clear difference in the *Bcl6* genotype comparing the two sorted populations (**Fig. 3B-C**). While around 30% of the non-Tfh cells had *Bcl6* indels, no such mutations were found in the Tfh population, in line with the essential role of *Bcl6* for Tfh differentiation. As we had established that the KRN+ iCR model was functional, we switched to targeting *Dnmt3a* as this gene is the most commonly somatically mutated gene in patients with rheumatoid arthritis (*16*). In contrast to *Bcl6*, we achieved a very high indel frequency in *Dnmt3a*, reflecting the sometimes stochastic nature of sgRNA activity. Interestingly, we found that *Dnmt3a* influenced the differentiation of autoreactive Tfh cells, as a significantly lower frequency of *Dnmt3a* indels was found in the Tfh cells compared to non-Tfh cells (**Fig. 3D-E**). In comparison, a difference was not found comparing sorted CD69+ and CD69-CD4+ T cells (**Fig. S3C**). This implied that *Dnmt3a* plays a positive role in Tfh differentiation. However, the role of *Dnmt3a* was notably weaker than the role of *Bcl6* in the differentiation.

**Figure 3.**
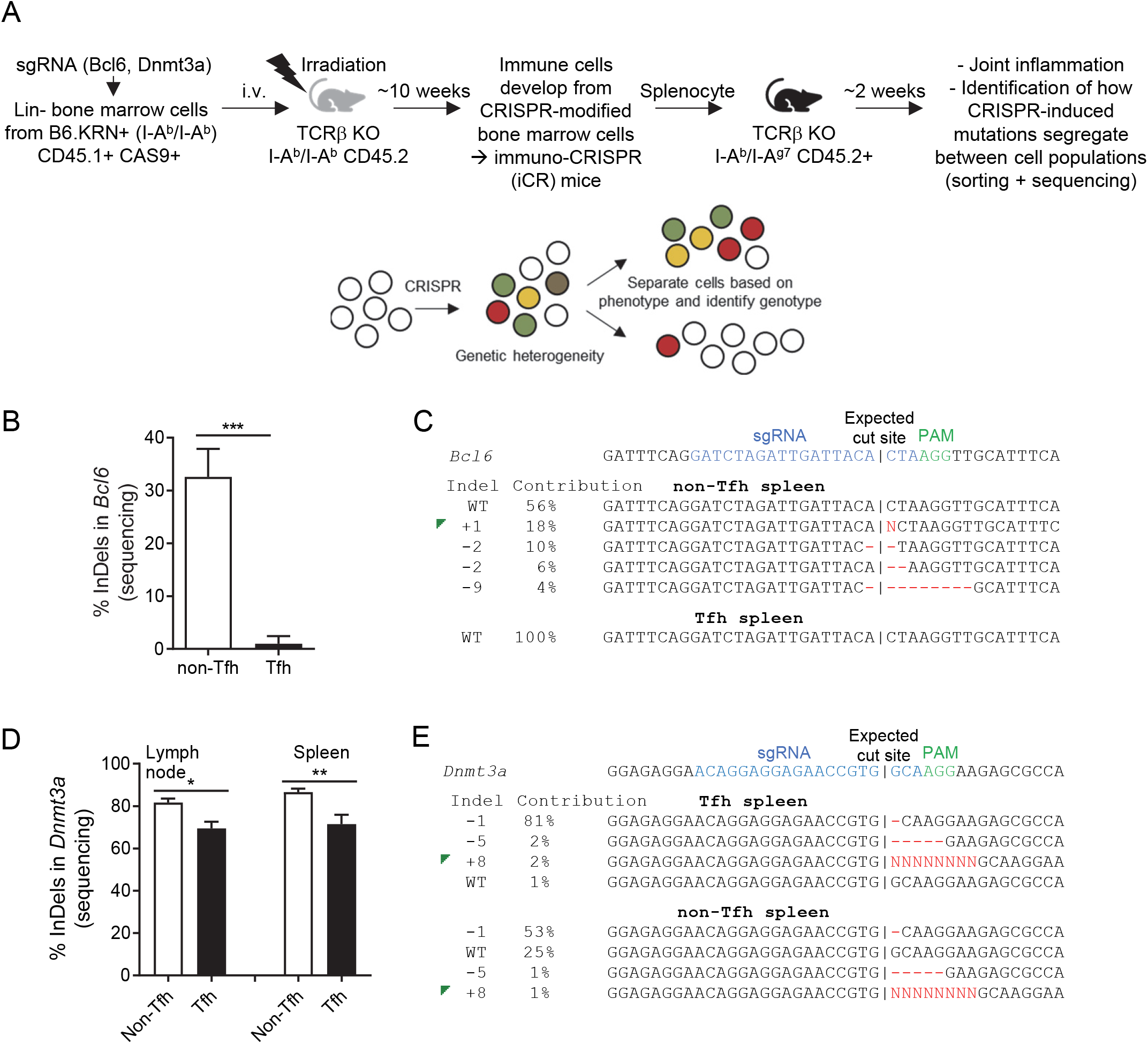
A CRISPR-based mixed bone marrow chimera identifies a role for Dnmt3a for autoreactive Tfh differentiation. (**A**) Schematics describing the experimental setup to generate immuno-CRISPR (iCR) mice with RISPR-induced genetic heterogeneity in the immune cell compartment, by transplanting CRISPR-modified cells from KRN+ mice (no disease as no I-A^g7^ is expressed). The modified cells are subsequently transferred to I-A^g7^+ recipient mice as in figure 1-2 resulting in joint inflammation. The cell intrinsic role of the targeted gene is reveled by sorting cell populations and comparing how mutations are segregating between populations (similar to a traditional mixed BM chimeric experiment). (**B**) *Bcl6* indel frequency in sorted splenic (viable, singlet, TCRb+, CD4+, PD1+, CXCR5+), and splenic non-Tfh T cells (viable, singlet, TCRb+, CD4+, R5-) isolated day 13 from TCRb KO I-Ab/I-Ag7 mice receiving splenocytes from KRN+ Bcl6 iCR mice. (**C**) Representative quantification of *Bcl6* genotype of one mice in (B). (**D**) *Dnmt3a* indel frequency in sorted splenic inguinal lymph node Tfh and non-Tfh cells isolated day 15 from TCRb KO I-Ab/I-Ag7 mice receiving splenocytes from KRN+ Dnmt3a iCR mice. (**E**) Representative quantification of *Dnmt3a* genotype of one mice in Data presented as mean and SEM (B, D), B n=5; D n=7-11. *** P <0.005, ** P <0.01, and * P <0.05 by ed T test (B, D). Data is representative for two or more independent experiments.

We concluded that the KRN+ iCR splenocyte transfer model could be used to study genes affecting the differentiation of autoreactive Tfh cells, and that *Dnmt3a* could be involved in promoting Tfh differentiation.

### Tfh differentiation is negatively impacted by the Dnmt3a^R878H^ mutation linked to clonal hematopoiesis

The iCR approach enables testing the function of a gene *in vivo* in a rapid and cost-effective manner and can help prioritize genes to study further. As we had identified a plausible role for *Dnmt3a*, we next breed CD4-cre mice with floxed *Dnmt3a* mice. The floxed *Dnmt3a* allele was generated so that when cre is expressed, a hotspot R878H mutation (replicating the human R882H mutation) linked to clonal hematopoiesis and AML is introduced (*26*). First, we generated a mixed BM chimera with WT CD45.1 BM and CD45.2 CD4-cre+ *Dnmt3a* fl/fl BM and immunized these with the T-cell dependent (TD) antigen sheep red blood cells (SRBCs) (**Fig. 4A**). In line with the iCR data, we found that a lower level of cells with the *Dnmt3a* alleles (CD45.2+) was found in the Tfh population compared to CD4+ non-Tfh T cells (**Fig. 4B**). This was evident in the spleen (**Fig. 4C**), typically studied in relation to i.v. SRBC immunizations, but interestingly, even more pronounced in Peyer’s patches (**Fig. 4D**).

**Figure 4.**
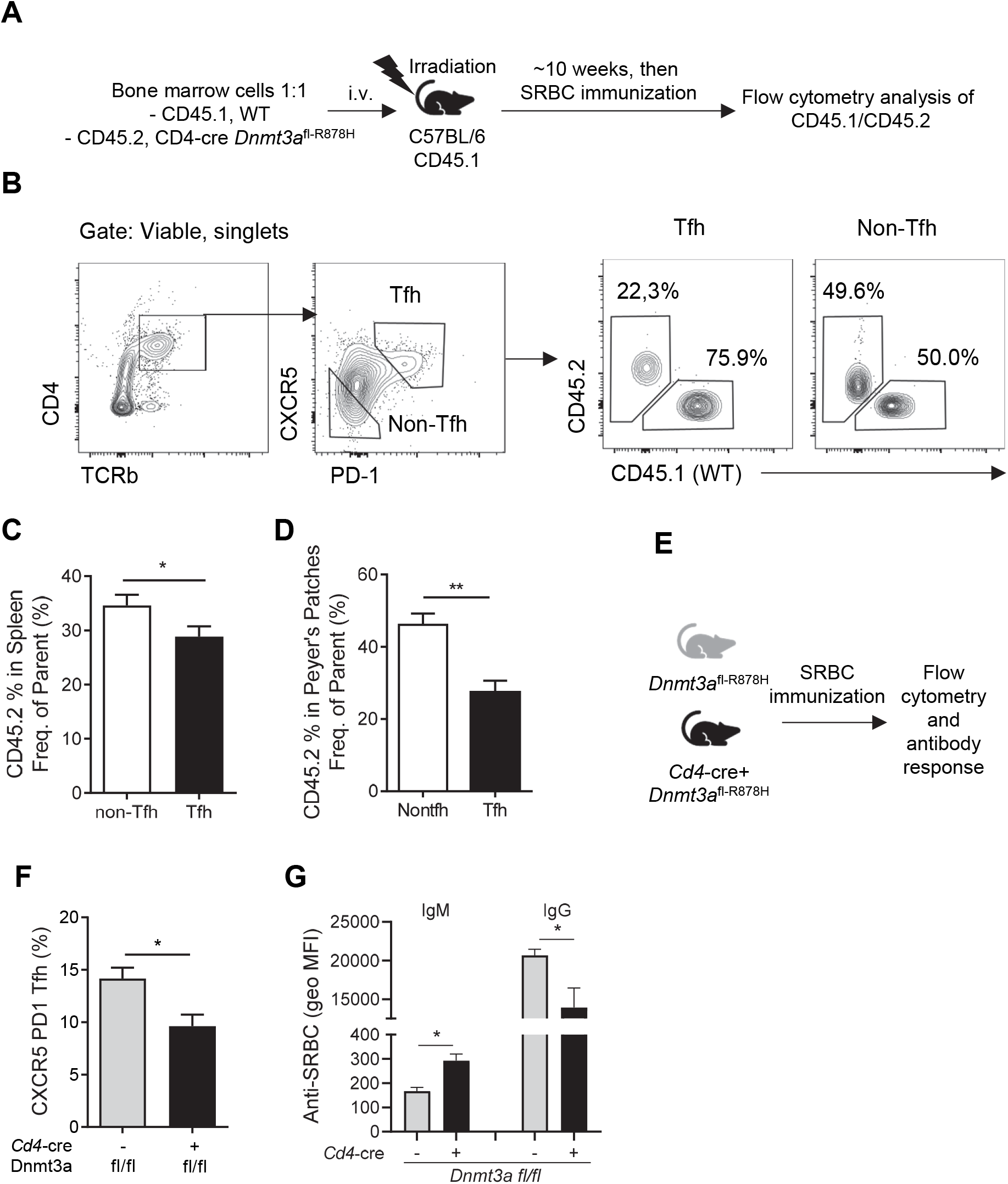
TFH differentiation is negatively impacted by the *Dnmt3a R878H* mutation linked to clonal hematopoiesis. (**A**) Schematics describing the setup of mixed BM chimera followed by SRBC immunization. (**B**)Representative FACS plots of gating of CD45.1 and CD45.2 Tfh cells in the mixed BM chimeras. (**C-D**) Frequency of cells with the *Dnmt3a R878H* mutation in splenic (C) and Payer’
ss patches (D) CD4+ Non-Tfh and Tfh cells. (**E**) Experimental setup SRBC immunization of *Dnmt3a R878H* (fl/fl) *+/-* CD4-Cre. (**F-G**) Frequency of splenic Tfh cells and analysis of antibody response (G) in mice with indicated genotype following SRBC immunization. body responses in (g) measured by flow cytometry as IgM and Data presented as mean and SEM (C, D, F G), C-D n=9; F-G n=3-5. *** P <0.005, ** P <0.01, and * P <0.05 by paired T test (C, D) and Mann-Whitney U-(F, G). Data is representative for two or more independent experiments.

Evaluating the development of immune cells in the mixed BM chimera showed that the B and T cell development was stable over time, while a somewhat unexpected increase in WT neutrophils was seen over time. We did not further analyze this phenotype, but the neutrophils are not supposed to be affected by the CD4-cre, although CD4-cre is known to not only delete in CD4+ T cells (also deleting in e.g. alveolar macrophages and all T cell subsets that pass through the thymic double positive stage (*27*)). We also observed that while CD44+ memory CD8+ T cells showed a 50/50 contribution of CD45.1/CD45.2 genotype, the CD44+ memory CD4+ T cells showed a clear bias with 70% being WT (CD45.1) (**Fig. S4**).

Finally, we immunized Dnmt3a fl/fl and CD4-cre Dnmt3a fl/fl mice with SRBC (**Fig. 4E**). In line with the iCR and mixed BM experiments, we found a lower frequency of Tfh cells in the floxed mice compared to the WT control (**Fig. 4F**).In addition, we found that 2 weeks after SRBC immunization, the CD4-cre Dnmt3a fl/fl group have higher anti-SRBC IgM level and lower anti-SRBC IgG antibody level compared with the CD4-wt Dnmt3a fl/fl group. (**Fig. 4G**) We concluded that the R878H *Dnmt3a* hotspot mutation negatively impacts Tfh differentiation during immunization with a standard T-dependent antigen.

## 4. Discussion

CD4+ T cells coordinate adaptive immune responses, including maladaptive autoimmune reactions, like in RA patients. The therapeutic efficacy of Abatacept, a drug that blocks the costimulatory interactions between activated T cells and antigen-presenting cells, shows that the activation of T cells plays an important role in disease initiation (*28*) and progression (*29*) of RA patients.

To enable detailed interrogation of autoreactive CD4+ T cells, we adopted the KRN+ splenocyte transfer model (*8*), a less commonly used version of the spontaneous K/BxN genetic model (*7*), and the K/BxN serum transfer model (*8, 30*) for autoreactive joint inflammation. A BM transfer version has also been established (*31*). Mechanistically, it has been well described by combining results from these models that when CD4+ KRN+ TCR transgenic T cells interact with I-Ag7+ B cells, the ensuing activation results in the production of IgG anti-GPI, which has arthritogenic activity. Notably, the T cell transfer model was characterized in more detail by LeBranche et al. (*22*) in 2010 but has not been widely adopted despite having several advantages (generating very reproducible joint inflammation which involves both an immunological priming and an effector phase of the disease).

Here, instead of transferring purified KRN+ T cells (*22*), we more crudely transferred KRN+ splenocytes, like in (*8*). The splenocyte transfer model, where cells from one mouse are given to 5-6 recipients, is much more experimentally straightforward than using purified T cells, generating more or less equivalent results in our experience. Still, there could be situations where a more controlled setup using purified T cells is preferred.

An initial characterization of the cellular dynamics following KRN+ splenocyte transfer (Fig. 1 and S1) showed that the transferred T cells (CD45.1+) expanded in several secondary lymphoid organs between day 3 and 6 after transfer. This correlated with the appearance of germinal center (GC) B cells stemming from the I-A^g7^+ B cells of the (CD45.2+) recipient mice. Concomitant with this, we found the production of IgM and IgG anti-GPI autoantibodies and a dramatic increase in joint inflammation between day 6 and 9. Interestingly, the joint inflammation day 0-12 correlated nicely with the appearance of IgG anti-GPI, in line with the expected autoantibody-driven etiology of the inflammation. However, as most cellular and serological parameters, including systemic IL-6 levels, declined day 15, the joint inflammation lingered, suggesting that both immunological processes and the resulting tissue response contribute to the clinical picture. An unexpected finding of the characterization was that the dynamics of CD69 expression on B cells and T cells were so different, with B cells peaking already day 3, i.e. before any evidence for clinical disease, and then declined, while T cell CD69 slowly increased expression with a peak at day 12. This could be compared to B and T cell CD69 upregulation following LPS injection, which shows a similar dynamic between B and T cells (*32*).

Of note, we typically don’t find many T cells in the inflamed joint, where cellular infiltrates are dominated by neutrophils (*18, 19*). Next, we wanted to confirm that the B-cell activation and subsequent production of autoantibodies were central to the pathology. To do this, we depleted B cells by injecting anti-CD20 every second day, starting 3 days before the splenocyte transfer (Fig. 2). In line with an essential role for B cells, no joint inflammation was seen in the anti-CD20 injected mice. Even more, transferring serum from mice with ongoing joint inflammation (triggered by KRN+ splenocyte transfer), showed that arthritogenic factors were present in serum (most likely autoantibodies, based on previous literature related to the KRN models (*8, 30*), and the B cell-depletion data).

As we had established the model’s premises, we wanted to test if we could apply the immuno-CRISPR (iCR) approach we previously developed (*23*). This approach is based on modifying BM cells with CRISPR-Cas9, targeting a gene of interest to generate a controlled genetically heterogeneous BM cell population. The modified BM cells are subsequently transplanted into an irradiated host, allowing the transplanted cells to develop into mature immune cells. As described in figure 3a, cells of interest could next be sorted, and the genetic heterogeneity compared between different cell populations to identify if the targeted gene affects the cell population. This approach shares many features with mixed BM chimeras, but uses sequencing instead of flow cytometry to quantify how different genotypes segregate between cell populations/phenotypes. Central to this approach is the fact that CRISPR-Cas9 doesn’t generate a homogenous KO population. Instead, a mixed genotype is typically seen, including mutations of different types as well as WT alleles. As we had established that the KRN+ T cells played a primary role in orchestrating the activation of autoreactive B cells, we decided to focus on Tfh differentiation. It’s well established that *Bcl6* is central to Tfh differentiation, and it has even been shown that BCL6 expression in KRN+ T cells is essential for pathology in one of the few studies using the KRN+ T cell transfer model (*25*). In line with that data, we found that Tfh differentiation in KRN+ *Bcl6* iCR cells completely depended on *Bcl6*. Thus, we could establish that the iCR approach was functional as a discovery approach in the KRN+ model. Next, we turned our focus to *Dnmt3a*. We had hypothesized that *Dnmt3a* mutations could contribute to autoreactive Tfh differentiation, but the data identified that *Dnmt3a*, in contrast, seemed to play the opposite role and that *Dnmt3a* mutations instead seemed to block Tfh differentiation partly. The phenotype was, however, not as profound as the *Bcl6* phenotype. Of note, we did not see any selective advantage of mutated *Dnmt3a* in cells comparing TCRb+ CD4+ CD69+ and TCRb+ CD4+ CD69-cells sorted from animals with ongoing joint inflammation, suggesting that the *Dnmt3a* mutations did not affect all features of CD4+ T cells.

The *Dnmt3a* iCR mice model identified that *Dnmt3a* could affect Tfh differentiation, and we decided to follow up the studies using more traditional animal models. To do this, we obtained *Dnmt3a* floxed R878H mice that, when crossed to a cre-line, introduce the R878H point mutation in exon 23, equivalent to one of the most common hotspot mutations found in individuals with clonal hematopoiesis (*26*). This mutation does not generate a null allele but vastly decreases the function of the gene (*33*). To target CD4+ T cells, we crossed the *Dnmt3a* floxed line with a CD4-cre. We first generated mixed BM chimeras with CD45.2+ cre+ *Dnmt3a* fl/fl BM and CD45.1+ WT BM cells and immunized these with SRBCs. In line with the iCR data, we found a preferential expansion of WT Tfh cells compared to non-Tfh cells, implying that *Dnmt3a* mutations negatively impacted Tfh differentiation also during a standard immunization protocol. Interestingly, this phenotype was the most pronounced in Peyer’s patches, where interactions with the microbiota trigger a continuous Tfh differentiation (*34*). Furthermore, two weeks after SRBC immunization, the CD4-cre Dnmt3a fl/fl group had significantly higher anti-SRBC IgM levels and lower anti-SRBC IgG antibody levels compared with the CD4-wt Dnmt3a fl/fl group, indicating that the Tfh phenotype affects have a distinct function.

## 5. Conclusion

Taken together, we can conclude that *Dnmt3a* mutations partly limit Tfh differentiation, which could negatively impact adaptive immune responses in affected individuals and thus potentially contribute to the suppressed immune responses seen in elderly individuals. Interestingly, recent literature identifies that *Tet2* deficiency can lead to the expansion of the Tfh population (*35-37*). In other words, the opposite phenotype to what we present here. This makes sense in light of the fact that DNMT3A and TET2 have somewhat different activities, with DNMT3A being a DNA methyl transferase, adding methyl groups, and TET2 is involved in removing DNA methyl groups.

## Acknowledgments

We are grateful to Eduardo Villablanca for providing mice, and to Taras Kreslavskiy for suggestions. This research was funded by grants from the Karolinska Institutet, Swedish Research Council, Swedish Cancer Society, Stiftelsen Professor Nanna Svartz Fond, and the China Scholarship Council. This work has additionally received support via the European Union/European Federation of Pharmaceutical Industries and Associations (EU/EFPIA) Innovative Medicines Initiative Joint Undertaking (RTCure Grant 777357).

FW has received consulting fees from SmartCella Solutions, which is outside the scope of the submitted study.

