## Supplementary material for "*Dnmt3a* mutations limit normal and autoreactive Tfh differentiation": Sup. Fig 1-4

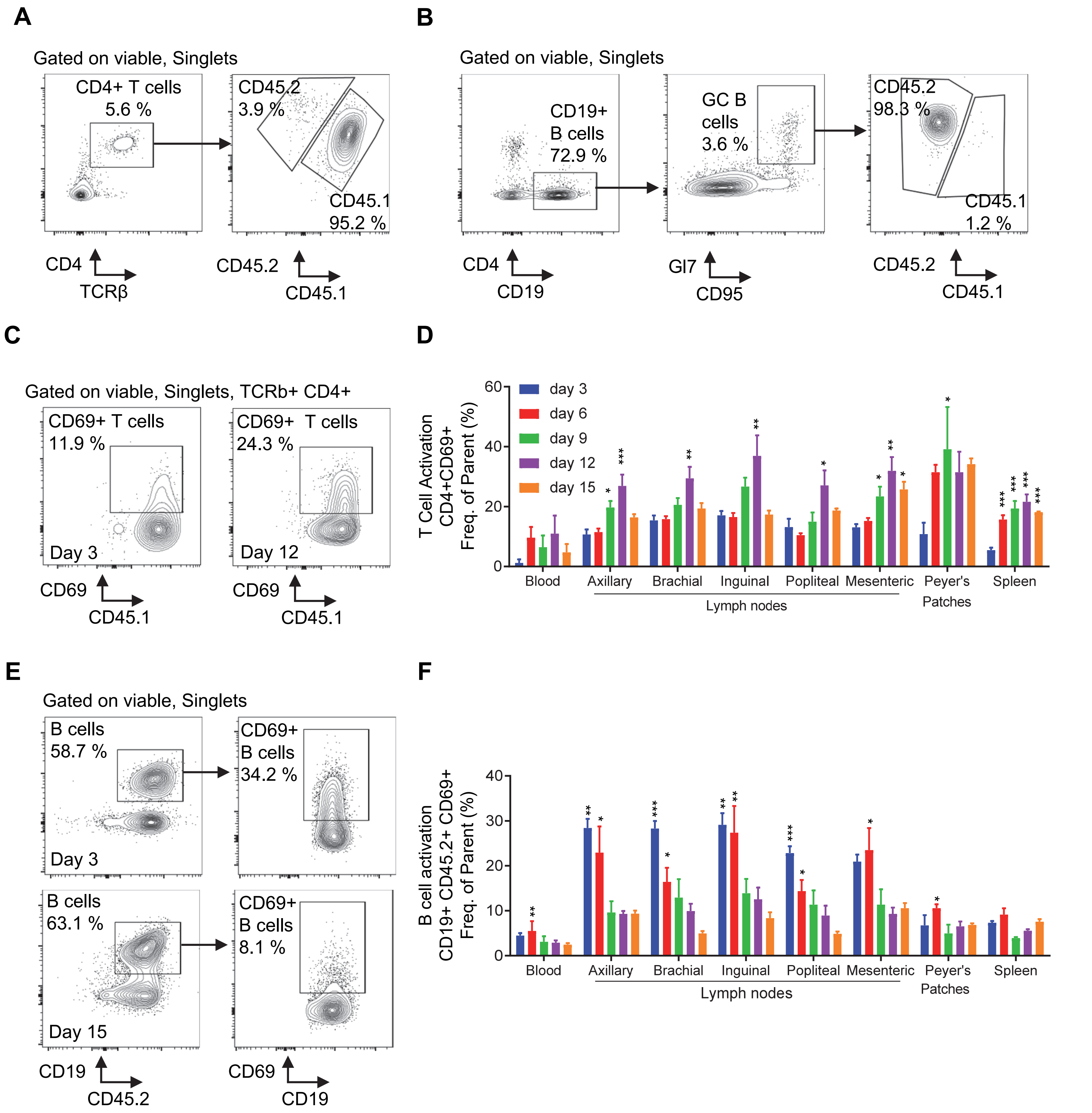


**Sup. Figure 1**. KRN+ splenocyte transfer results in a reproducible and timed joint inflammation. (**A-B**) Representative gating for flow cytometry of CD4+ T (A), GC B cells (B), (**C-D**) Frequency of CD69+ T cells (of total CD45.1+ CD4+ T cells) detected by flow cytometry (C shows examples from day 3 and day 9) in different organs at indicated timepoints (D). (**E-F**) Frequency of CD69+ B cells (of total CD45.2+ CD19+ B cells) detected by flow cytometry (E shows examples from day 3 and day 15) in different organs at indicated timepoints (F). Data presented as mean and SEM (D and F, n = 3-4). *** P <0.005, ** P <0.01, and * P <0.05 by one-way ANOVA with Dunnett´s post hoc test comparing different timepoints to the day 3 group (D), or the day 15 group (F). Data is representative for two or more independent experiments.

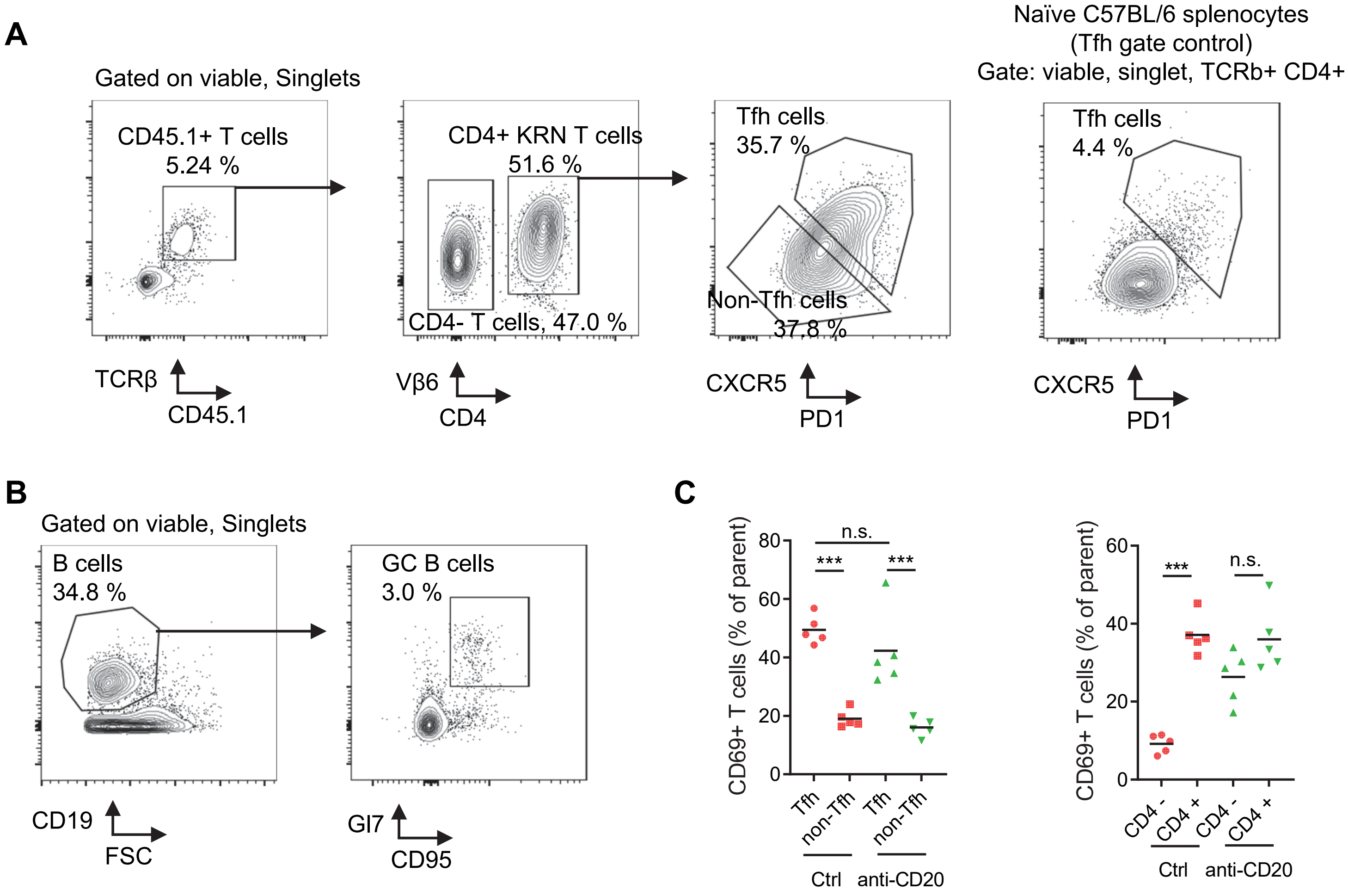


**Sup. Figure 2.** Joint inflammation in the KRN T cell transfer model is abrogated by B cell depletion. (**A-B**) Representative gating for flow cytometry in fig 2. (**C**) Quantification of splenic CD69+ T cells day 9, separated based on CD45.1+ TCRb+ CD4+ Tfh and Cd45.1+ TCRb+ CD4+ non-Tfh (left) or CD45.1+ TCRb+ CD4+ and CD4- cells (right). . *** P <0.005, and ns = non-significant by one-way ANOVA with Tukey's post hoc test.

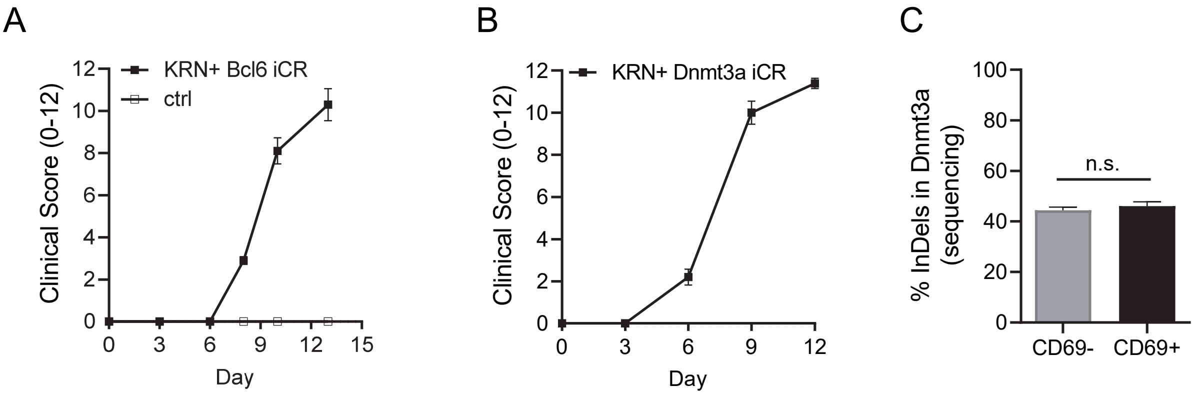


**Sup. Figure 3.** A CRISPR-based mixed bone marrow chimera identifies a role for Dnmt3a for autoreactive Tfh differentiation. (**A-B**) Disease development in mice receiving KRN+ splenocytes with CRISPR-induced *Bcl6* (A) and *Dnmt3a* (B) mutations. (**C**) *Dnmt3a* indel frequency in sorted splenic CD69+ and CD69- CD4+ T cells isolated day 12 from TCRb KO I-Ab/I-Ag7 mice receiving splenocytes from KRN+ Dnmt3a iCR mice. ns = non-significant by paired T test (C). Data is representative for two or more independent experiments.

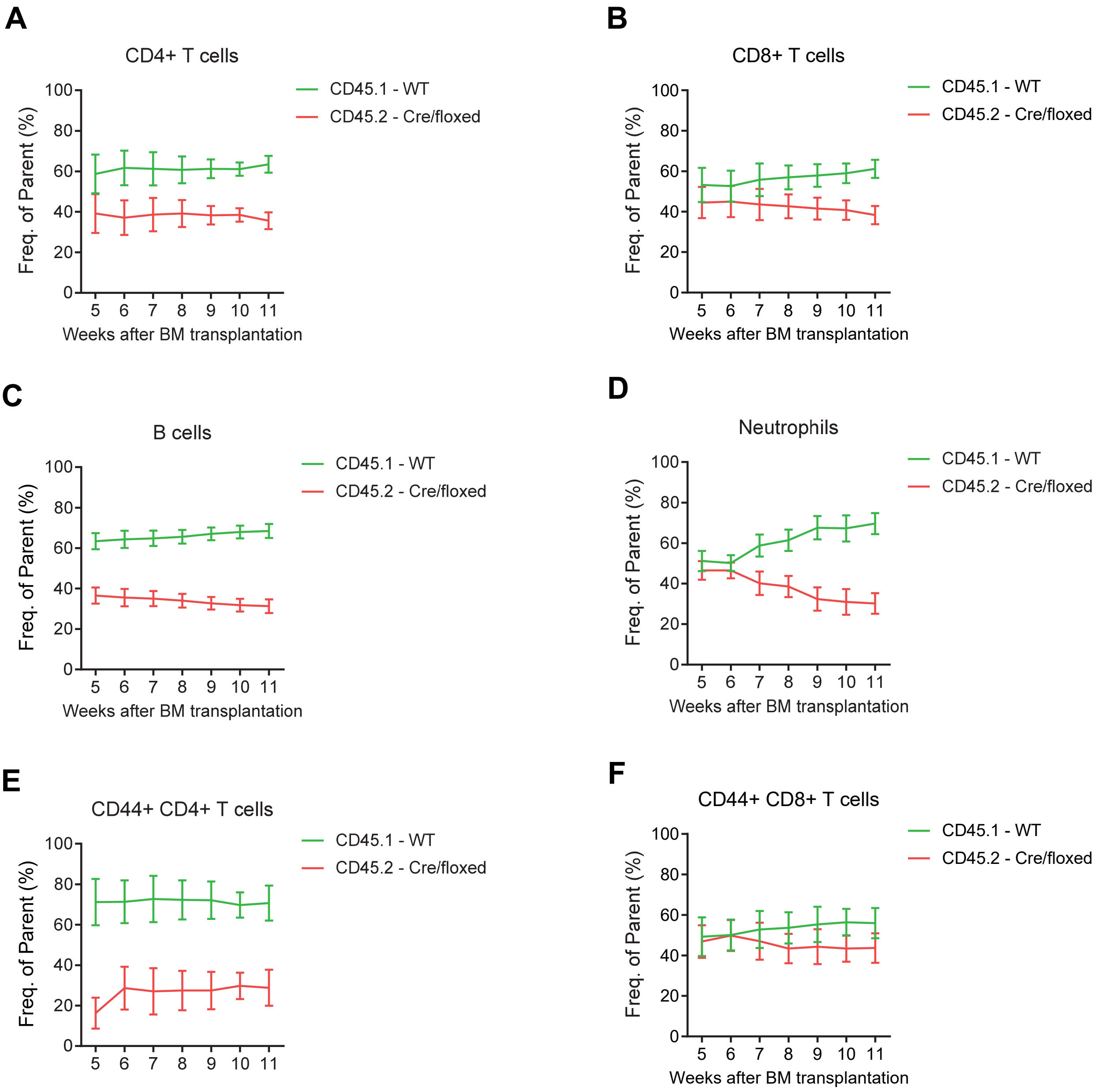


**Sup. Figure 4**. TFH differentiation is negatively impacted by the Dnmt3a R878H mutation linked to clonal hematopoiesis. (**A-F**) Contribution of the two mixed bone marrows to the indicated cell population over time (n=9).

### Supplemental Tables

Table S1 Genotyping primers

| Primer name | Sequence (5' – 3') |
|---|---|
| *KRN TCR alpha chain Fw* | AGGTCCACAGCTCCTTCTGA |
| *KRN TCR alpha chain Rv* | GTATTGGAAGGGGCCAGAG |
| *KRN TCR beta chain Fw* | GGGCAAAAACTGACCTTGAA |
| *KRN TCR beta chain Rv* | GAGCCTGGTTGTTTGTGGAT |
| *I-A$^{g7}$ Fw* | TTCAAGGGCGAGTGCTACTT |
| *I-A$^{g7}$ Rv* | GTTCGCTCCAGGTACTGCTT |
| *I-A$^{b}$ Fw* | TTCATGGGCGAGTGCTACTT |
| *I-A$^{b}$ Rv* | CGTTCGCTCCAGGATCTC |
| *Dnmt3a Fw-com* | CTCCTTGGATTTGAGGAGGA |
| *Dnmt3a Rv-wt* | TGCACATGAGAACTGGATGG |
| *Dnmt3a Rv-mut* | ATTAAGGGCCAGCTCATTCC |
| *CD4$^{cre}$ com-chr3* | AACTTGCACAGCTCAGAATGC |
| *CD4$^{cre}$ mut-chr3* | TTAGGGTGGGGCTCAGAAGG |
| *CD4$^{cre}$ wt-chr3* | ACCTGAGATTCCACCAAACTTGA |

Table S2 sgRNA sequences used in the study

| Gene | Sequence (5' – 3') |
|---|---|
| *Bcl6* | UAGUGUAAUCAAUCUAGAUC |
| *Dnmt3a* | ACAGGAGGAGAACCGUGGCA |

Table S3 primers used to detect sgRNA targeted region

| Primer name | Sequence (5' – 3') |
|---|---|
| *Bcl6-Fw* | TGTGTCGACAACATGCTCCA |
| *Bcl6-Rv* | AGTCTGCTAGGCTTTCCACG |
| *Dnmt3a-Fw* | GGCCAACGCTTGGAATTGAA |
| *Dnmt3a-Rv* | GTGTCCTCTCTCACATGACCG |

Table S4 Antibody List

| Antibody | Company | Ref. No. | Clone |
|---|---|---|---|
| CD45.1-BV605 | Biolegend | 110738 | A20 |
| B220-APC/Cy7 | BD Bioscience | 552094 | RA3-6B2 |
| B220-PE | Invitrogen | 12-0452-82 | RA3-6B2 |
| CD11b-PerCP/Cy5.5 | Biolegend | 101228 | M1/70 |
| CD19-AF647 | Biolegend | 115522 | 6D5 |
| CD3-APC | Invitrogen | 17-0032-82 | 17A2 |
| CD44-PE/Cy7 | Biolegend | 103030 | IM7 |
| CD45.1-APC/Cy7 | BD Bioscience | 560579 | A20 |
| CD45.1-BV605 | BD Bioscience | 110738 | A20 |
| CD45.1-FITC | BD Bioscience | 561871 | A20 |
| CD45.2-BV785 | Biolegend | 109839 | 104 |
| CD45.2-BV785 | Biolegend | 109839 | 104 |
| CD4-FITC | Invitrogen | 11-0042-85 | RM4-5 |
